# The UV potentiated mutational signature of clinical stage G-quadruplex binder CX5461

**DOI:** 10.64898/2026.04.10.717570

**Authors:** Elena Zaikova, Damian Yap, Armaghan Sarvar, Auden Hafezi, Jay Tan, Viviana Cerda, Kaylee Li, Daniel Lai, Karen Gelmon, John Hilton, Lesley Seymour, Peter Stirling, Dave Cescon, Samuel Aparicio

## Abstract

Drug-related UVA-induced photoreactions have been reported for several therapeutic compounds, including fluoroquinolones. CX5461 is a clinically relevant quinolone-derived anti-cancer small molecule with documented UVA-sensitising activity. Here we compared, by bulk and clonal whole genome sequencing (WGS) under light-protected conditions, the mutational signatures in human retinal pigment epithelial cells (RPE1) exposed to UVA, CX5461, or co-exposed to UVA and CX5461. Treatment with CX5461 or UVA alone resulted in a low SNV burden and background-like mutational profiles. In contrast, bulk sequencing of human cells co-exposed to UVA and CX5461 had a markedly higher SNV burden characterized by T>A and T>C substitutions. Furthermore, single-cell clonal expansion and sequencing of CX5461 alone, UVA alone or CX5461+UVA treatments confirmed that the pattern was only observed when cells were exposed to both UVA and CX5461. The CX5461+UVA-associated SNV signature we report arises only when CX5461-treated cells are exposed to UVA, and is not observed when CX5461-treated cells are shielded from light. We do not observe strong single base mutagenic activity of CX5461 alone, under light protected conditions. Our data emphasise the need for appropriate controls and light-exposure precautions when studying base mutagenesis activity of known photosensitiser molecules.

## Introduction

We and others have discovered that small molecule compounds which stabilize DNA secondary structures from stacking of guanine bases (G-quadruplexes, G4) induce DNA damage and transcription-replication conflicts^5–7^. In the backgrounds of DNA repair-deficient cancers, principally homologous recombination repair deficiency (HRD), but also non-homologous end joining (NHEJ) and TMEJ deficiency^5,8–10^, G4-ligands are selectively active. G4 stabilising small molecules, such as CX5461 and pyridostatin, induce double-stranded breaks in epithelial cells by at least two mechanisms (polymerase stalling^11^, helicase and topoisomerase interference^10,12^), have strong selective activity with loss of specific DNA repair functions including HRD, non-homologous end joining (NHEJ), polymerase-theta (TMEJ) and ubiquitin-mediated DNA damage response (DDR; genes RNF168, UBE2N)^8^. Genetic hypersensitivity has also been observed with genes mediating transcription and chromatin regulator functions, such as SMARCA4, ATRX and DDX36^8,9^. The mechanism of selectivity involves at least two factors. First, the formation of more G4 folded regions in cancers^13^, which occurs due to duplication of DNA and increased open chromatin regions. Second, the inability of cancers with repair or chromatin mutations to either repair or tolerate the mutational burden arising from G4 stabilization results in cell death^5,11^, in contrast to normal tissues where these functions are intact. This leads to a clinical hypothesis that tumours where critical chromatin, transcription or DNA repair functions are inactivated, will have a therapeutic window for G-quadruplex small molecule binders.

We recently tested this therapeutic hypothesis in patients with a phase 1 trial of CX5461, a molecule we discovered to be a potent G4-ligand^11^. We observed^4^ evidence of clinical activity on-target in participants with cancers related to germline-BRCA2 and PALB2 mutations, validating the conceptual approach. The molecule tested was broadly well tolerated at doses that also resulted in significant clinical responses, notably without dose-dependent anaemia or other chemotherapeutic end-organ effects. Beyond G4 stabilizing activity, CX5461 was also noted in two clinical trials (ANZCTR 1261300106172; NCT02719977 also referred to as IND231) to be a potent UV photosensitizer^3,4^. Fluoroquinolones are well-established UVA photosensitizers, which can lead to increased DNA damage, in part through the formation of cyclobutane pyrimidine dimers when exposed to UV light^4^. The UV photosensitization tendency is a class effect associated with the quinolone structure and not a general feature of G4 ligands^4^. Although G4 DNA damage effects of CX5461 have been reported in cell lines and model organisms, the UV mutation spectrum of CX5461 requires further clarification. Here we report the single and double base substitution and indel signatures of CX5461 in human RPE1 cells exposed to UV. Joint UV+CX5461 exposure resulted in a bulk WGS detectable mutational burden, with a distinct mutational pattern characterized by T>A, T>C mutations.

## Results

### The CX5461+ UVA mutational signature in human cells is distinct from CX5461 or UV exposure alone

To characterize the distribution and types of short variants induced by CX5461 photosensitization, we analyzed CX5461 and UV co-exposure *in vitro*, with deep bulk WGS and single cell clone expansion protocols (Figure 1a). First, we treated retinal pigment epithelial cell lines (RPE1) Cas9 wildtype (WT) or p53 knockout (TP53) cells (Supplementary Table 1) with CX5461 alone, with UV alone, or the combination of both CX5461 and a metered dose of UVA (Supplementary Table 2). Precautions were taken to shield CX5461-only conditions from stray light exposure (foil screening, avoidance of incubators with UV bulbs and using only indirect illumination when manipulating plates). The concentrations of drug, UV and combination were chosen to ensure at least 50% of cells survived the most cell toxic condition (CX drug + UV). Treated and untreated control cells were subjected to deep whole genome sequencing (median 75X for bulk and 40X for expanded clones, Supplementary Table 2), and treatment-associated mutations were identified using the untreated cells as the matched normal. Notably, in WGS bulk-sequenced samples, treatment with UV alone, or CX5461 alone, resulted in fewer than 150 detected single base substitutions (SBS) per condition, whereas the combination of CX5461 and UV resulted in a 2-4 orders of magnitude increase in SBS mutations (Supplementary Table 2) and had a distinct mutational pattern characterized by T>N (Supplementary Figure 1b). Since bulk WGS may miss low frequency distributed single-base mutagenic activity, we also validated the signatures observed by serial treatment, using single cell clonal expansion and WGS of clones, to capture any fixed mutations. We again took care to shield the non-UV drug only conditions from incident light and UV. The mutational patterns present in the cloned cells were characterised by the same computational pipeline. We again observed the same T>N mutational pattern in UV+CX5461 with a high burden of SBS (76744, 117326 median in WT and TP53), contrasted with at least tenfold-lower mutational abundance and only background mutational signatures in UV alone (1594, 6729 median in WT and TP53) or CX5461 alone (2329, 6107 median in WT and TP53) (Supplementary Figure 1b, Supplementary Table 2). To examine the possible origin and similarity of these single nucleotide substitutions, we calculated the cosine distance of the trinucleotide context of each treatment condition (Figure 1b). This revealed that bulk-sequenced and cloned cell SBS profiles were near identical (cosine similarity 0.95-1) between all UV+CX samples with a 50mJ/cm^2^ dose (Figure 1b). We extracted two signatures from the bulk and two signatures from the cloned-cell RPE1 datasets. The cloned cell and bulk signature sets were near-identical, where one of the signatures was a UV+CX signature, and the other represented the background (Figure 1c). Signature refitting using non-linear least squares optimization (NNLS) showed that the identified signatures represented almost the entire observed mutational profile in RPE1 cells (Supplementary Figure 1c and 1d). The observed SBS-UVCX signature is dominated by T>A; T>C and T>G trinucleotide contexts (Figure 1c), and is reminiscent of a recently reported psoralen induced mutation signature^1^ (Figure 2, cosine similarity ~0.67), as well as the previously reported single base signature of UV+CX5461 exposure^14^ in *C. elegans* (cosine similarity 0.67-0.73). Notably, the SBS-UVCX signature is near-identical to that of Koh et al^15^ (cosine distance 0.96-0.99, Figure 1d), who described this pattern as a mutational signature of unknown origin from CX5461 exposure in RPE1 cells. In contrast, RPE1 samples treated with either UV or CX5461 alone, exhibit lower numbers of SBS with a predominant C>A, C>T trinucleotide context, and similar SBS signatures (cosine similarity ~0.85) to SBS40. This SBS40-like signature is highly similar to the “Background” signature reported by Koh et al^15^ (cosine similarity 0.91-0.98, Figure 1d). The UV and CX-only treated samples additionally show similarity to SBS signatures that are ascribed to background, clock-like (SBS5) and exposure to tobacco (SBS4, SBS92, SBS29) or azathioprine patterns (Figure 2).

**Figure 1.**
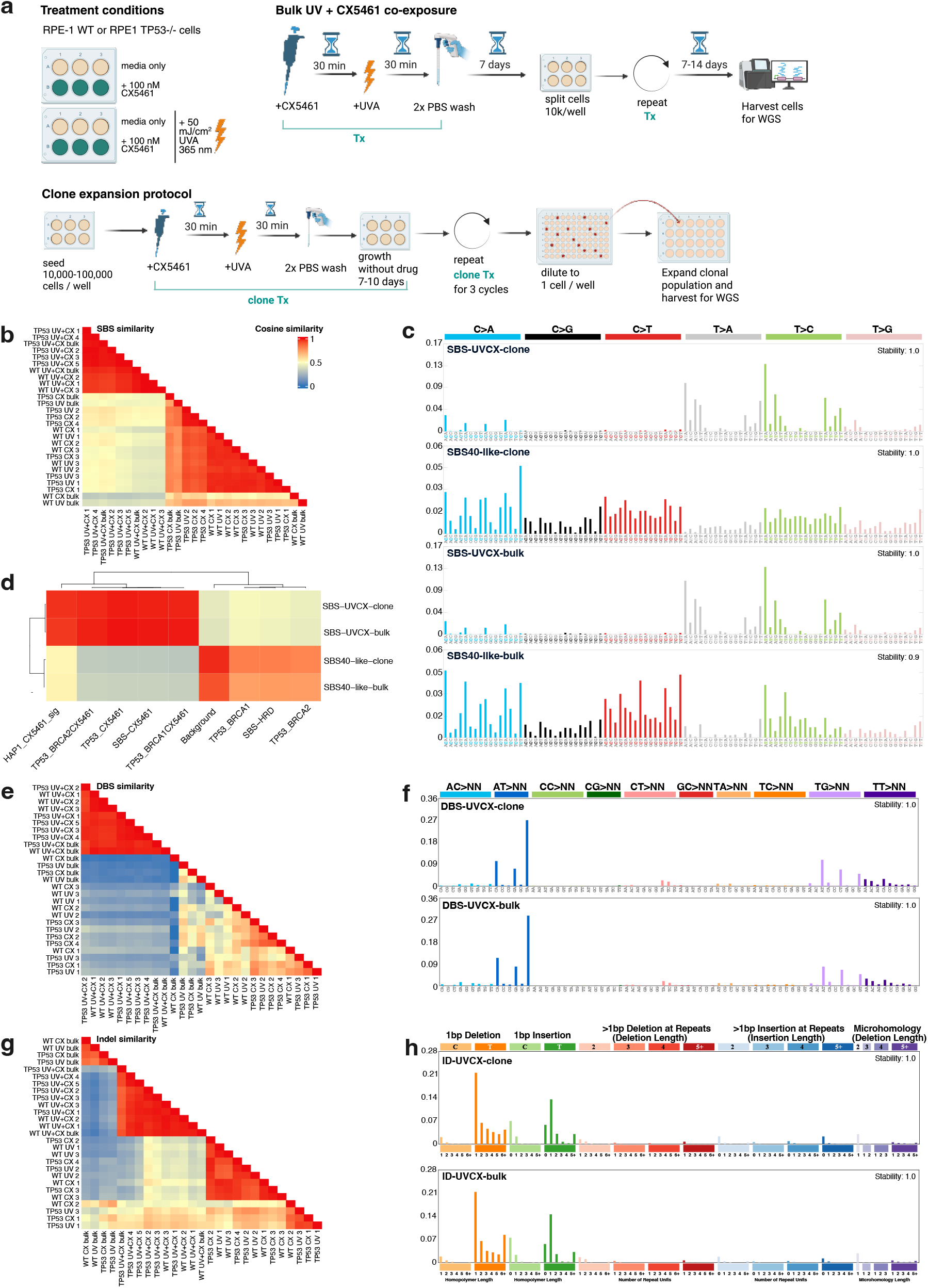
Mutational signatures of immortalized RPE1-hTERT Cas9 (WT) and RPE1-hTERT Cas9 p53KO (TP53) cells. (**a**) Experimental design and treatment details of bulk-sequenced and single clone RPE1 samples. To minimize exposure to ambient UVA, plates were covered in foil for all incubation steps. The schematic was generated using BioRender (https://app.biorender.com/). (**b**) Cosine similarity of SBS mutational profile of WT and TP53 RPE1 cells, treated with 50 mJ/cm^2^ UVA and/or CX5461. Sample axis labeled as: genotype+treatment bulk or 1,2,3 for clones. (**c**) SBS signatures extracted from clone-expanded and bulk-sequenced RPE1 cells, exposed to UVA and/or CX5461. (**d**) Cosine similarity of SBS signatures extracted from WT and RPE1 cells with SBS signatures reported by Koh et al^15^. Cosine similarity of (**e**) DBS and (**g**) Indel mutational profiles in all CX5461, UVA or co-exposed RPE1 cell line data. Sample axis labeled as: genotype+treatment bulk or 1,2,3 for clones. DBS (**f**) and indel (**h**) signatures associated with UVA and CX5461 co-exposure identified in clone and bulk-sequenced RPE1 datasets.

**Figure 2.**
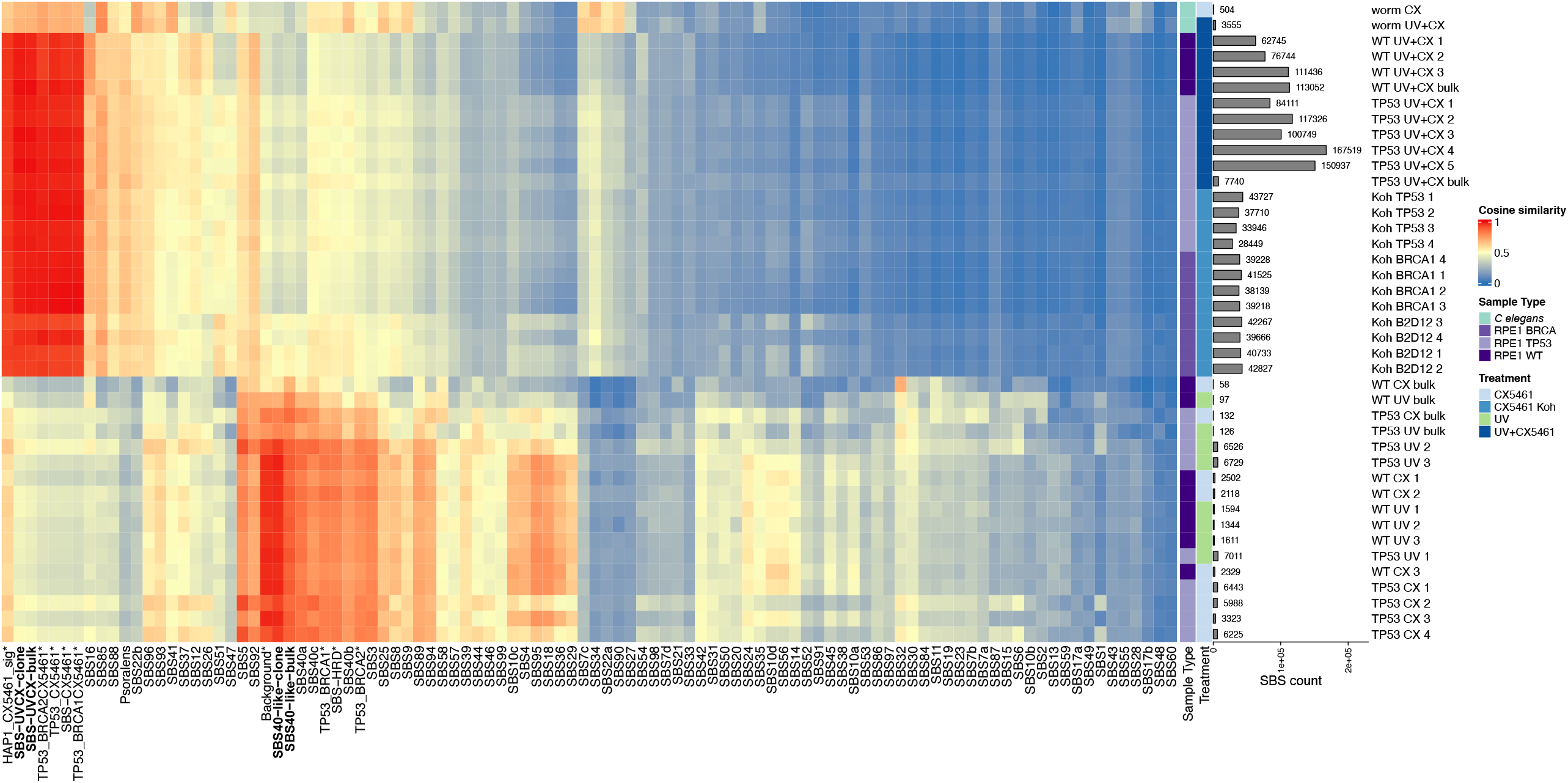
Cosine similarity between mutational profiles of *C. elegans* data, bulk-sequenced and clone expansion RPE1 cell data as well as CX5461-treated RPE1 samples from Koh et al and SBS signatures, including COSMIC v3.4, SBS signatures from Koh et al, indicated with an asterisk and the signatures identified in our RPE1 data, in bold font.

Although less is known about the assignment of dinucleotide (Figure 1e) and indel (Figure 1g) mutational patterns, we identified UV+CX signatures in both mutation types (Figure 1f and 1h). The DBS-UVCX signature is characterized by AT>NN, TG>NN and TT>NN substitutions (Figure 1f), accompanied by a 10-to 100-fold increase in DBS mutation abundance (Figure 3b). We also identified a background DBS signature in the clone samples, but not the bulk (Figure 3a). Neither signature is highly similar to any COSMIC DBS signatures (Figure 3c). Similarly, indels also show a characteristic mutational pattern accompanied by a 10-to 100-fold increase in mutations under CX+UV (Supplementary Figure 2c). Although the bulk and clone ID-UVCX signatures are near-identical (Figure 1g and 1h), the background indel signatures identified in clone and bulk are different (Supplementary Figure 2a). The ID-UVCX is somewhat similar (cosine similarity 0.7-0.73) with ID23, which is attributed to damage involving bulky lesions such as aristolochic acid exposure (Supplementary Figure 2b). The clone background ID signature is somewhat similar to ID2, associated with slippage during DNA replication (cosine similarity 0.76, Supplementary Figure 2b). This is consistent with CX5461 with UV producing UV-induced CPD bulk adducts which require nucleotide excision repair (NER)^4^. As observed with SBS and DBS, the indel profiles of RPE1 samples from Koh et al^15^ are highly similar to our UV+CX co-treated cells (median cosine similarity 0.90), but not with UV or CX5461 (Supplementary Figure 2e). We note that the UV potentiated SNV (SBS, DBS and ID) signatures of CX5461 observed here occurred at a dose of 50 mJ/cm^2^ delivered over 16 seconds (irradiance = 3.125 mW/cm^2^) ~2 orders of magnitude below the intensity used for phototherapy and similar to UVA irradiance on a sunny day (mean ~5.4mW/cm^2^) at mid-latitude^16^, where 1 hour of sunlight would deliver ~19,440 mJ/cm2, ~400x greater dose and ~9 secs exposure would yield the 50mJ/cm^2^ dose used here. This emphasises the strong UV photosensitization properties of CX5461.

**Figure 3.**
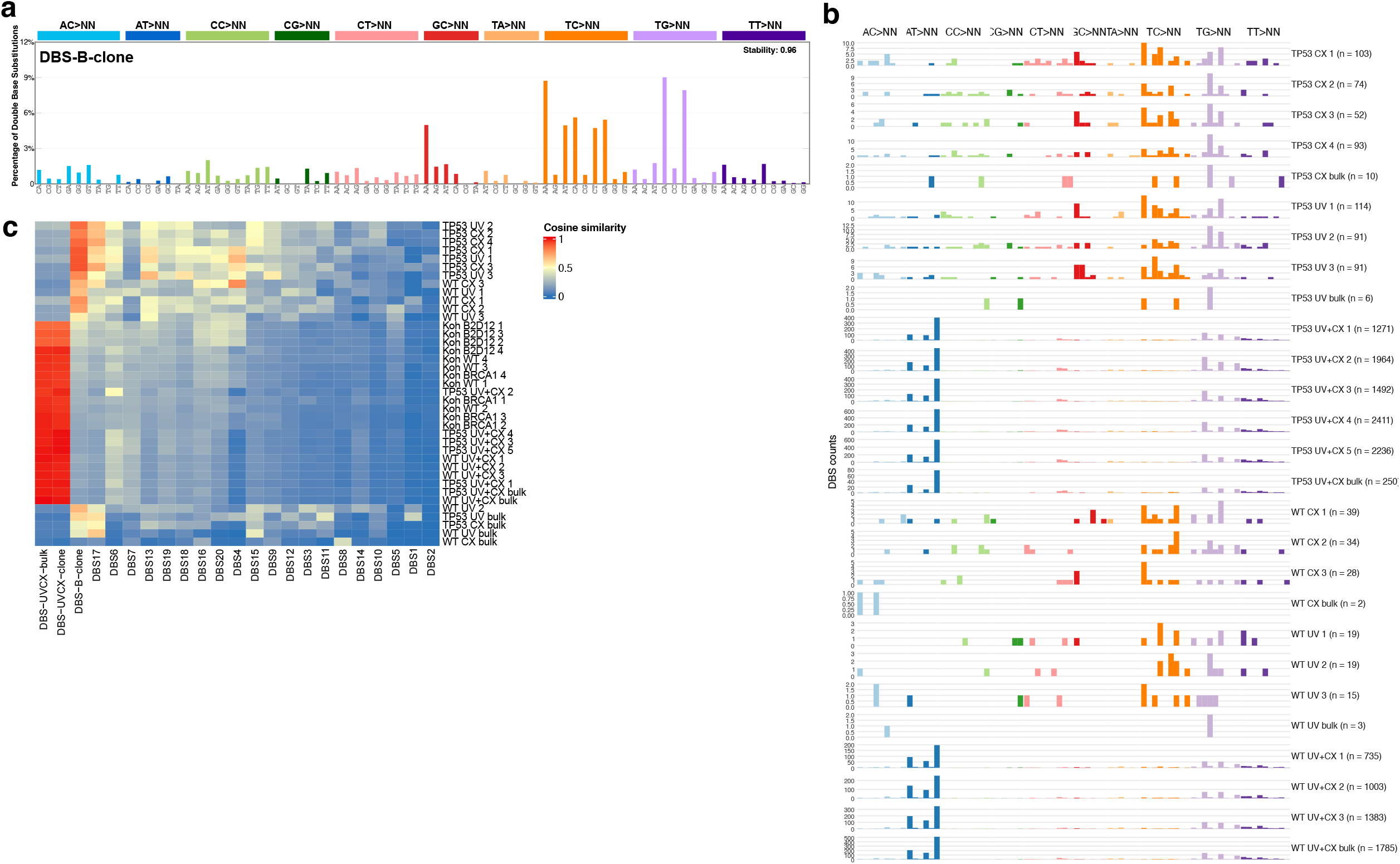
Distribution of DBS mutations in RPE1 cell data. (**a**) Proportion of 78 DBS mutation types in the second DBS signature identified in the clone expansion data. (**b**) Distribution of DBS mutations in CX5461-treated, UVA-treated or co-exposed bulk-sequenced RPE1 TP53 and WT cells and all clone expansion replicates. (**c**) Cosine similarity between mutational profiles of bulk-sequenced and clone expansion RPE1 cell data, CX5461-treated RPE1 samples from Koh et al^15^ and DBS signatures identified in our RPE1 data as well as those from COSMIC v3.4.

## Discussion

Clastogenic agents and base mutagens are used widely in oncology, however monitoring of residual mutational effects in somatic tissues is not normally analysed as part of clinical trials. CX5461 is known to create mutations through at least two processes. First, single and double stranded breaks followed by error prone repair or absence of homology directed repair leads to small indels and large scale deletions, partially enriched at G-quadruplex sites ^6^. This relates to a core mechanism of action as a DNA repair deficiency selective small molecule. A second process is through the highly potent UV photosensitization induced by CX5461, which has been previously documented *in vitro*^*4*^, as an SBS signature in *C. elegans*^*14*^ and as a clinically manifest avoidable toxicity^3,4^. This activity is a class effect of the CX5461 molecule, which shares quinolone structural features with other known photosensitizers, and is not a general property of G-quadruplex binders. Although the photosensitization effect of CX5461 has been clearly documented in earlier clinical and pre-clinical studies and the mutational signature reported in *C. elegans*, the human cell mutational spectrum of UV+CX5461 potentiation has not been clearly documented. Here we characterized the single nucleotide mutational signature of UVA and CX5461 co-exposure in a human retinal pigment epithelial cell type that is commonly used for genomic mutagenesis studies. Human retinal pigment epithelial cells exposed deliberately to short pulses of UVA+CX5461 exhibited dramatic increases in detectable SNVs, with a distinct T>N trinucleotide context. The UV+CX signature is readily detectable in both bulk sequencing experiments and in single-cell cloning experiments, which are more sensitive for detecting low frequency base mutations. We note the signature of UV+CX in our experiments is nearly identical to that of an earlier report from Koh et al in the same cell line type, using the same single cell cloning method, but attributed as previously unsuspected massive base mutagenic activity of CX5461. This earlier report does not mention control experimental data nor explicit screening precautions for possible stray UV exposure which is common in laboratory environments, pertinent to working with a known highly potent photosensitizer. Nor was the previously described *C*.*elegans* UV+CX5461 signature analysed in context and as we show here this also bears similarity. Critically, when light screening precautions are taken, exposure of RPE cells with CX5461 alone, in our hands, does not exhibit the reported Koh et al SBS activity and only mutational processes that are associated with background processes or sequencing artefacts were detected. We conclude that the previously reported “unanticipated massive collateral single-nucleotide mutagenic activity” attributed to CX5461, can best be interpreted as that due to UVA+CX5461 co-exposure mutational signature rather than intrinsic mutagenicity of CX5461 alone. While our work was in review, a preliminary report of duplex DNA sequencing from CX5461 exposed human tissue samples of an earlier phase 1 trial noted no large excess of SBS mutations in 4 patients followed serially^17^. This is consistent with an absence of intrinsic base mutagenic activity in the absence of direct UVA exposure in the sampled tissues. CX5461 (compound 7c in ^18^) was also previously reported as non-mutagenic in an Ames assay. Photosensitization is a well documented clinical side effect of CX5461, avoidable with the photoprotection measures recommended for ongoing human studies. Our report documents the UV+CX5461 SBS signature in human cells and highlights the need for *in vitro* light screening precautions and controls when assaying single base mutagenesis of known photosensitizers.

## Materials and Methods

### Cell lines

Immortalized hTERT RPE1 cells were sourced from ATCC and transduced with the Lenti-FLAG-Cas9-2A-Blast vector and selected with blasticidin. Single cell clones were assayed for Cas9 expression, which was confirmed by immunoblotting^19^. RPE1 cas9 p53 KO cells were generated using hTERT RPE1 expressing Cas9 and sgRNAs targeting TP53 (CAGAATGCAAGAAGCCCAGA) to generate the CRISPR knockouts^20^. These batches of cells were obtained from the Stirling Lab (BC Cancer Research Institute, Vancouver, BC, Canada). Both lines of RPE1 Cas9 cells were authenticated via STR profiling (Genetica Cell Line Testing, Burlington, NC, USA using the Promega PowerPlex 16 HS kit), which showed that the transduced cells retained the parental RPE1 profile (ATCC CRL-4000) at all tested loci (Supplementary Table 1).

### Drugs

CX-5461 for cell line experiments was purchased from Selleckchem (Houston, TX, USA) and dissolved in 50 mM NaH_2_PO_4_ and stored in 1 mM aliquots with protection from light.

### Drug and UV treatment of RPE1 cells

RPE1-hTERT Cas9 (WT) and RPE1-hTERT Cas9 p53KO (TP53) grown in DMEM/F-12 or Advanced DMEM/F-12 (Gibco), supplemented by 10% fetal bovine serum (FBS), were split at 10,000 cell per well of a 6-well plate. The following day, 50-100 nM CX5461 was added in DMEM/F-12 +10% FBS (without phenol-red) to the cells and incubated for 30 min before being irradiated at 50 mJ/cm^2^ UVA (UVP Crosslinker CL-1000L, 365nm, Analytik Jena, Upland, CA, USA) in the plate without the lid before being replaced in the incubator for a further 30 min. The cells were then washed twice with PBS and then grown in DMEM/F-12 or Advanced DMEM/F-12 supplemented with 10% FBS without any drug for a further 5-7 days or until confluent (Figure 1a). For bulk sequencing, cells were split at 10,000 cells per well and subject to a second round of UVA irradiation. Cells were allowed to regrow (2-3 weeks) and then harvested for bulk whole genome sequencing (Supplementary Figure 1a). Cells used for clone expansion were split at 100,000 cells per well and subjected to a second and third round of UVA irradiation with the drug as before (Figure 1a). Cells were allowed to regrow for 2-3 weeks, and were subsequently diluted. Only wells with a visually confirmed single cell were selected for clonal outgrowth and expansion. Clones were then harvested for DNA extraction and whole genome sequencing. Flow cytometric cell sorting (FACS) was avoided due to the probability of exposing the cells to laser light.

For all experiments, plates were covered in aluminum foil when not being manipulated (Supplementary Figure 1a) to reduce exposure to stray sources of UV. Where possible, care was taken to avoid exposure to light by working in the hood without lights (and relying only on indirect illumination in the room) when the plates are manipulated without foil covering. Incubators without UV sterilization bulbs were used. The 50–100 nM CX-5461 range corresponds to the clinically measured mean trough concentrations (~50 ng/mL or ~0.1 µM^4^, which persist for several days between weekly doses, i.e. a sub-therapeutic level of exposure.

### Whole Genome Sequencing

Genomic DNA from cell pellets were obtained using the Wizard® Genomic DNA Purification Kit (Promega, Madison, WI, USA) following the manufacturer’s protocol. Libraries were constructed using the Illumina PCR-Free Genome Library Protocol (Genome Science Center, Vancouver, BC, Canada) and sequenced to an average depth of 80X or 40X for samples following clone expansion.

### Sequence processing, short variant calling and mutational signature comparison

Whole genome sequencing reads were aligned to the hg38 reference genome using bwa-mem 0.7.15-r1140^21^. Following alignment, technical artifacts were corrected by marking duplicates and recalibrating base quality using GATK best practices workflows (version 4.1.3.0)^22,23^. Short variants, including indels and single-base, dinucleotide, and trinucleotide mutations, were called using Mutect2 (version 2.2) in matched normal mode^24^. Untreated control cell line samples and skin punch biopsies collected prior to CX5461 treatment were designated as the matched normal samples for short variant calling. Candidate variants were subsequently filtered to remove likely germline variants, contamination and artifacts using FilterMutectCall and FilterAlignmentArtifacts (version 4.1.3.0). Variants that passed filters were then used for mutational signatures and distribution analyses. Whole genome sequencing data for RPE1 cell lines from Koh et al^15^ were processed using the high stringency pipeline described above. Untreated replicate samples for each genotype were combined into a single untreated “matched normal” alignment file. Subsequent variant calling for each CX5461-treated replicate was performed using the combined sequence data for the corresponding RPE1 genotype. We compared our filtered variant calls with those reported by Koh et al^15^. We report a higher number of variants, as we did not filter based on minimum VAF, we recovered ~95% previously reported with highly similar mutational patterns. All replicates were highly similar in mutational profiles and burden, and had high cosine similarity.

We used non-negative matrix factorization (NMF) for *de novo* extraction of SBS, DBS and indel signatures from the mutational profiles of RPE1 bulk and clonal expansion samples using SigProfilerMatrixGenerator^25^ and SigProfilerExtractor^26^. A maximum of 10 signatures were evaluated, with 100 NMF replicates, with the most stable solution selected for final signatures. To compare the mutational profiles with known mutational signatures, we downloaded COSMIC SBS, indel and DBS signatures (v3.4, GRCh38 reference) from https://cancer.sanger.ac.uk/signatures/downloads/, and added signatures relevant to UV sensitization^1,15^ to this reference set of signatures. The relative contribution of known signatures in the mutational profiles of our samples, we used non-linear least squares refitting. Statistical analyses and data visualization were performed using R 4.3.2, and the MutationalPatterns^28^, ggplot2^29^, BSgenome and vcfR R packages.

## Data availability

Whole genome sequencing alignments and variant calls are available in EGA under EGAS50000001145.

## Acknowledgements

S.A. holds the Nan and Lorraine Robertson Chair in Breast Cancer and is a Canada Research Chair in Molecular Oncology (950-230610). Additional funding was provided by the Terry Fox Research Institute Grant 1082, Canadian Cancer Society Research Institute Impact program Grant 705617, CIHR Grant FDN-148429, Breast Cancer Research Foundation award (BCRF-24-180), Canada Foundation for Innovation (40044) to S.A.

## Author contributions

EZ, DY, SAp, designed the study, JH, KG, LS, DC led the IND231, E.Z., A.S., D.L., A.H., J.T. K.L. performed genomic and statistical analyses. D.Y. and V.C. carried out cell line drug treatment and sequencing experiments. S.A., E.Z., D.Y. and A.S. wrote/edited the manuscript. All authors reviewed and approved the manuscript.

## Figure Legends

**Supplementary Figure 1.**
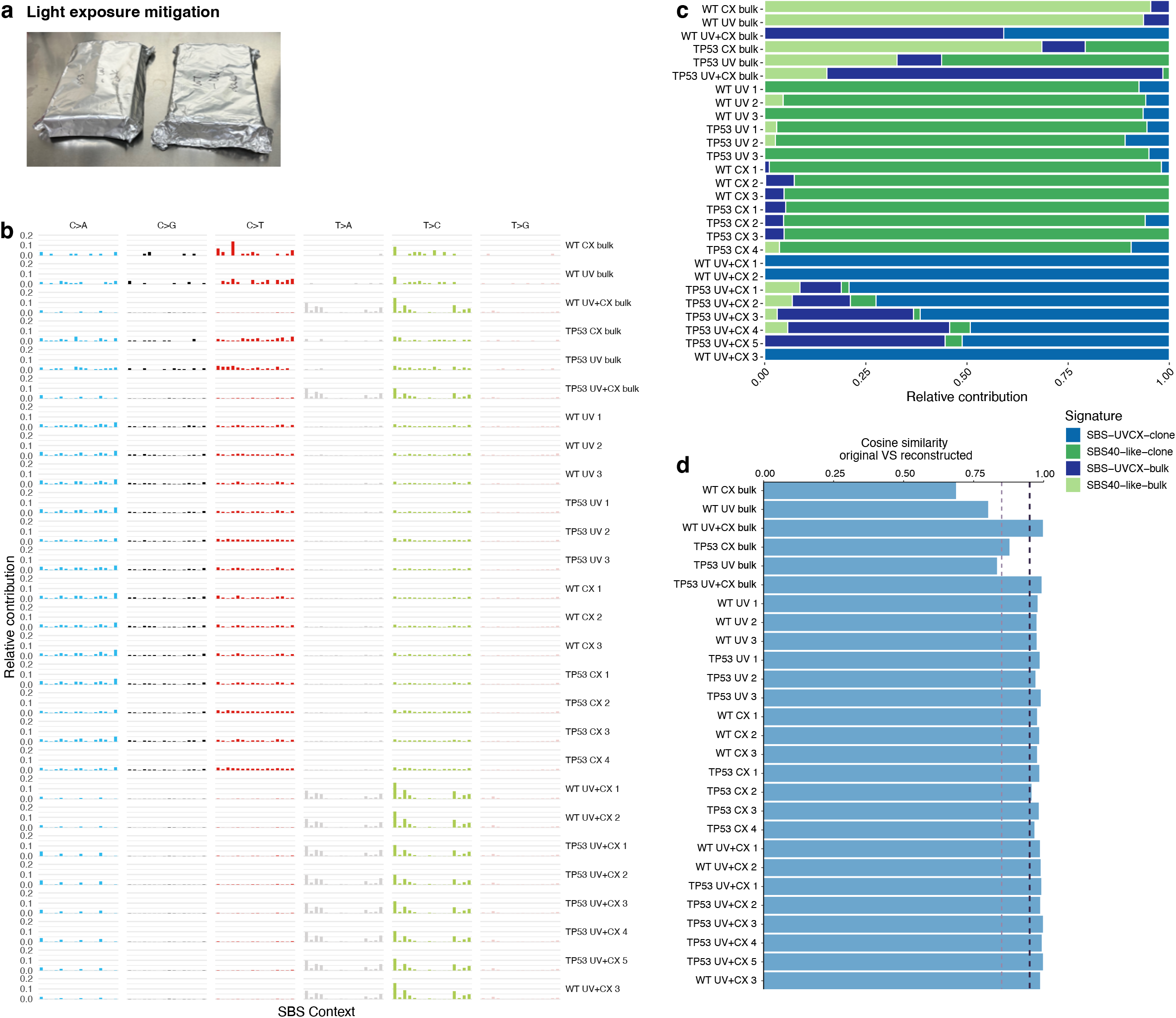
RPE1 SBS mutational profiles across treatments. (**a**) Image of cell culture plates wrapped in aluminum foil for mitigation of exposure to light. (**b**) Distribution of SBS mutations in CX5461-treated, UVA-treated or co-exposed RPE1 TP53 and WT cells, including all clone expansion replicates and bulk sequenced samples. (**c**) Relative contribution of the signatures, as determined by non-linear least squares refitting, identified in the RPE1 cell line data across CX5461-treated, UVA-treated or co-exposed RPE1 TP53 and WT samples. (**d**) Cosine similarity of original and reconstructed matrix based on the four signatures we identified in our RPE1 data. The dashed lines represent cosine similarity of 0.85 (light purple) and 0.95 (dark blue).

**Supplementary Figure 2.**
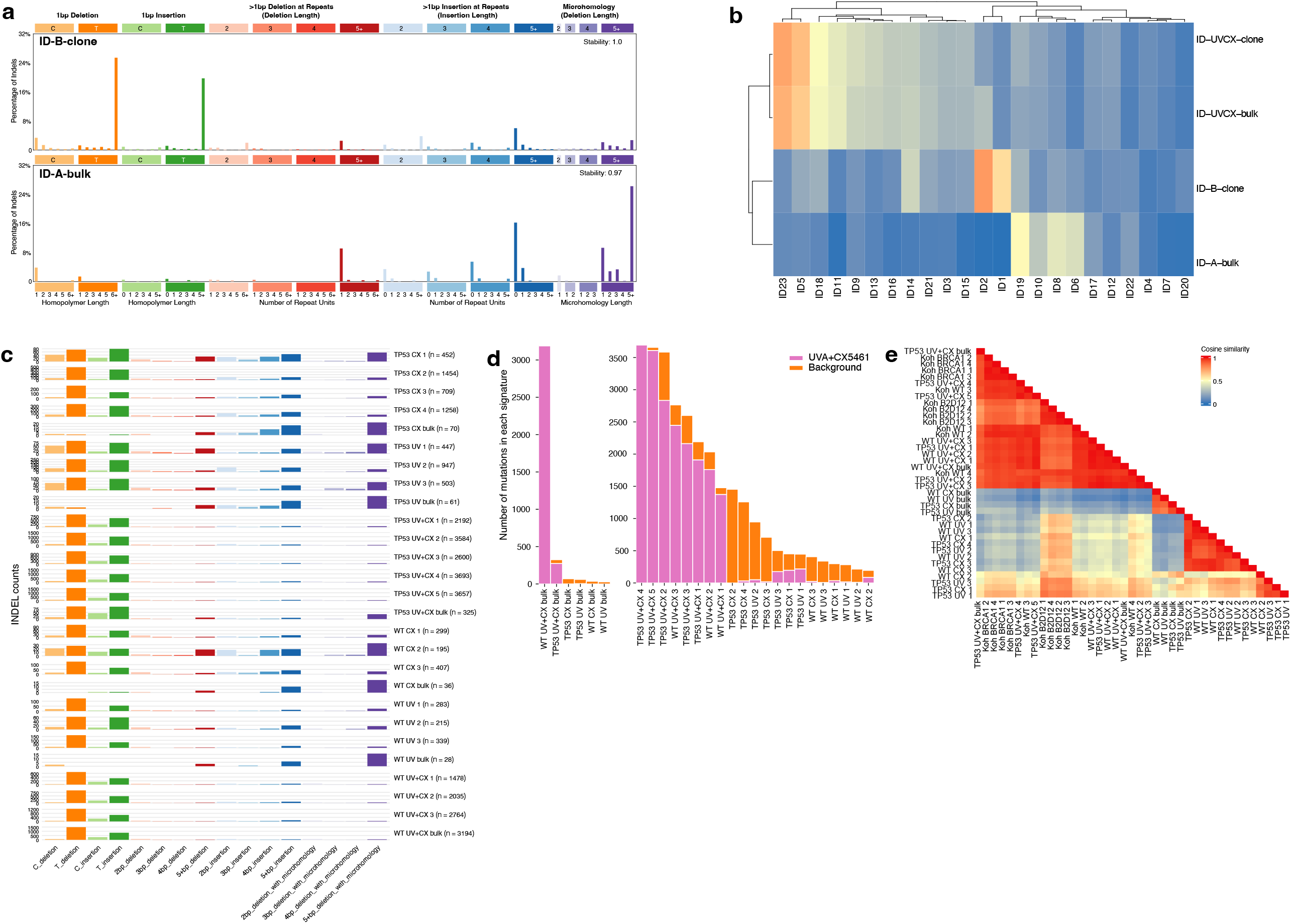
Distribution of indel mutations in RPE1 cell data. (**a**) Proportion of 83 indel mutation types in the second indel signatures identified in clone expansion data (top) and bulk-sequenced RPE1 data (bottom). (**b**) Cosine similarity of the indel signatures extracted from our data and COSMIC v 3.4 indel signatures. (**c**) Distribution of indel mutation types and indel counts in bulk sequenced WT and TP53 RPE1 cells and all clone expansion replicates. (**d**) Signature activity (UV+CX signatures and the secondary/background signature) in RPE1 cell line treated with UVA and/or CX5461. (**e**) Cosine similarity of indel mutation distributions between treated bulk and clone RPE1 cells in the present study and CX5461-treated RPE1 samples from Koh et al^15^.

## Supplementary material

**Supplementary Table 1.** Cell line identity confirmation by Short Tandem Repeat profiling.

| STR profiling | hTERT RPE1 Cas9<br>WT | hTERT RPE1 Cas9<br>TP53 KO | hTERT RPE1 (ATCC CRL-4000) |
| --- | --- | --- | --- |
| Amelogenin | X | X | X |
| CSF1PO | 12,14 | 12,14 | 12,14 |
| D13S317 | 11,12 | 11,12 | 11,12 |
| D16S539 | 11 | 11 | 11 |
| D5S818 | 11 | 11 | 11 |
| D7S820 | 10,11 | 10,11 | 10,11 |
| TH01 | 9 | 9 | 9 |
| TPOX | 8 | 8 | 8 |
| VWA | 17,18 | 17,18 | 17,18 |
| Penta_D | 9,11 | 9,11 | 9,11 |
| Penta_E | 12,14 | 12,14 | 12,14 |
| D3S1358 | 15,16 | 15,16 | 15,16 |
| D21S11 | 31,32.2 | 31,32.2 | 31,32.2 |
| D18S51 | 14,16 | 14,16 | 14,16 |
| D8S1179 | 10 | 10 | 10 |
| FGA | 20,23 | 20,23 | 20,23 |
| D19S433 | n.d. | n.d. | 14,16.2 |
| D2S1338 | n.d. | n.d. | 19,24 |
| Mycoplasma | Negative | Negative | Not detected |

**Supplementary Table 2.** Sequencing coverage and the number of acquired mutations in sequencing data from RPE1 cell lines.

| Sample | Treatment | Mean coverage | Coverage sd | Number of SBS | Number of DBS | Number of indels |
| --- | --- | --- | --- | --- | --- | --- |
| TP53 Control bulk | Untreated control | 82.04 | 23.49 | - | - | - |
| TP53 CX bulk | 100nM CX5461 | 87.22 | 25.67 | 132 | 10 | 70 |
| TP53 UV bulk | 50mJ/cm<sup>2</sup> UVA | 69.83 | 21.08 | 126 | 6 | 61 |
| TP53 UV+CX bulk | 50mJ/cm<sup>2</sup> UVA + 50nM CX5461 | 74.39 | 21.15 | 7740 | 250 | 325 |
| TP53 Control 1 | Untreated control | 44.12 | 13.63 | - | - | - |
| TP53 Control 2 | Untreated control | 37.65 | 10.74 | - | - | - |
| TP53 CX 1 | 50nM CX5461 | 37.08 | 10.99 | 6443 | 103 | 452 |
| TP53 CX 2 | 50nM CX5461 | 67.98 | 18.22 | 5988 | 74 | 1454 |
| TP53 CX 3 | 50nM CX5461 | 45.78 | 13.09 | 3323 | 52 | 709 |
| TP53 CX 4 | 50nM CX5461 | 39.78 | 11.68 | 6225 | 93 | 1258 |
| TP53 UV 1 | 50mJ/cm<sup>2</sup> UVA | 42.97 | 12.44 | 7011 | 114 | 447 |
| TP53 UV 2 | 50mJ/cm<sup>2</sup> UVA | 41.62 | 13.17 | 6526 | 91 | 947 |
| TP53 UV 3 | 50mJ/cm<sup>2</sup> UVA | 45.77 | 14.10 | 6729 | 91 | 503 |
| TP53 UV+CX 1 | 50mJ/cm<sup>2</sup> UVA + 50nM CX5461 | 42.74 | 12.75 | 84111 | 1271 | 2192 |
| TP53 UV+CX 2 | 50mJ/cm<sup>2</sup> UVA + 50nM CX5461 | 34.49 | 10.05 | 117326 | 1964 | 3584 |
| TP53 UV+CX 3 | 50mJ/cm<sup>2</sup> UVA + 50nM CX5461 | 36.55 | 10.38 | 100749 | 1492 | 2600 |
| TP53 UV+CX 4 | 50mJ/cm<sup>2</sup> UVA + 50nM CX5461 | 38.11 | 11.07 | 167519 | 2411 | 3693 |
| TP53 UV+CX 5 | 50mJ/cm<sup>2</sup> UVA + 50nM CX5461 | 41.14 | 12.14 | 150937 | 2236 | 3657 |
| WT Control bulk | Untreated control | 87.90 | 22.54 | - | - | - |
| WT CX bulk | 100nM CX5461 | 69.17 | 18.10 | 58 | 2 | 36 |
| WT UV bulk | 50mJ/cm<sup>2</sup> UVA | 78.50 | 20.80 | 97 | 3 | 28 |
| WT UV+CX bulk | 50mJ/cm<sup>2</sup> UVA + 50nM CX5461 | 90.10 | 22.03 | 113052 | 1785 | 3194 |
| WT Control 1 | Untreated control | 56.07 | 14.68 | - | - | - |
| WT Control 2 | Untreated control | 44.63 | 12.23 | - | - | - |
| WT CX 1 | 50nM CX5461 | 35.00 | 10.26 | 2502 | 39 | 299 |
| WT CX 2 | 50nM CX5461 | 37.61 | 10.85 | 2118 | 34 | 195 |
| WT CX 3 | 50nM CX5461 | 40.40 | 11.57 | 2329 | 28 | 407 |
| WT UV 1 | 50mJ/cm<sup>2</sup> UVA | 40.18 | 11.28 | 1594 | 19 | 283 |
| WT UV 2 | 50mJ/cm<sup>2</sup> UVA | 36.63 | 10.53 | 1344 | 19 | 215 |
| WT UV 3 | 50mJ/cm<sup>2</sup> UVA | 44.49 | 12.22 | 1611 | 15 | 339 |
| WT UV+CX 1 | 50mJ/cm<sup>2</sup> UVA + 50nM CX5461 | 39.94 | 11.32 | 62745 | 735 | 1478 |
| WT UV+CX 2 | 50mJ/cm<sup>2</sup> UVA + 50nM CX5461 | 39.66 | 11.45 | 76744 | 1003 | 2035 |
| WT UV+CX 3 | 50mJ/cm<sup>2</sup> UVA + 50nM CX5461 | 48.49 | 13.18 | 111436 | 1383 | 2764 |

